# CoffeeProt: An online tool for correlation and functional enrichment of proteome-wide systems genetics

**DOI:** 10.1101/2020.10.02.323246

**Authors:** Jeffrey Molendijk, Marcus M. Seldin, Benjamin L. Parker

## Abstract

The integration of genomics, proteomics and phenotypic traits across genetically diverse populations is a powerful approach to discover novel biological regulators. The increasing volume of complex data require new and easy-to-use tools accessible to a variety of scientists for the discovery and visualization of functionally relevant associations. To meet this requirement, we developed *CoffeeProt*, an open-source tool that analyzes genetic variants associated to protein networks and phenotypic traits. *CoffeeProt* uses proteomics data to perform correlation network analysis and annotates protein-protein interactions and subcellular localizations. It then integrates genetic and phenotypic associations along with variant effect predictions. We demonstrate its utility with the analysis of mouse and human population data enabling the rapid identification of genetic variants associated with protein complexes and clinical traits. We expect *CoffeeProt* will serve the proteomics and systems genetics communities, leading to the discovery of novel biologically relevant associations. *CoffeeProt* is available at www.coffeeprot.com.

## INTRODUCTION

The field of genetics has realized significant progress in the discovery of phenotype-associated genetic variation in recent years (1). As of 2018, over 4,000 genes harboring variants causal for monogenic traits have been discovered in addition to over 60,000 significant genome-wide associations across many human diseases and traits. This success is attributable to technological advances, access to increasing amounts of genetic and phenotypic data, and the continuous development of novel analytical tools. The functional relevance of phenotype-genotype associations in diverse populations and environments is a rapidly evolving area contributing to our understanding of complex health and disease traits. Systems genetics is the approach in which intermediate molecular phenotypes are examined in relation to genetic variation to improve our understanding of complex traits and common diseases (2). Evidently, the integration of different biological layers has distinct advantages over the analysis of a single biological layer as the flow of information can be modelled to identify and prioritize core regulators for functional validation (3). Mapping genetic loci to complex traits via quantitative trait loci (QTL) analysis is a powerful approach of combining biological layers. Such traits could include proteins (pQTL), transcripts (eQTL) and other molecular traits (molQTL) (4). However, a significant challenge of these studies is that the number of associations can be very large, requiring complex integrations, filtering and data visualizations to interpret biologically relevant interactions and discover potential new causal relationships for subsequent validation. ProGem is a recent tool developed by Stacey *et al.* to identify and prioritize causal genes at molecular QTLs (5). This powerful framework leverages positional and eQTL data combined with pathway analysis to prioritize possible genes underlying the biological mechanisms. Additionally, Seldin *et al.* collated a list of computational resources used in the field of systems genetics (6) including the statistical and modelling tools Mergeomics (7), WGCNA (8), MEGENA (9), MOFA (10) and ARACNE (11). These tools are used to identify key drivers in biological pathways, constructing gene co-expression networks and inferring relationships between quantitative measures. However, a key limitation of many of these tools is they either require advanced experience in computational analysis, are not available online or they do not have visualization features. This means they may not be accessible to scientists with a variety of systems genetic skill levels and the time required to learn these computational protocols may be prohibitive. Furthermore, many of these tools are not focused on the inclusion of proteomic data with additional annotations such as proteinprotein interactions or subcellular localization. The inclusion of this data offers exciting opportunities to further investigate the mechanisms of genotype and trait associations. For example, the integration of pQTLs with protein-protein interaction network analysis has the capacity to define genetic regulation of protein complexes (12,13) or the inclusion of co-regulated network analysis with subcellular localization and molQTLs can identify compartmentalized associations (14). However, the annotation of large volumes of proteomics and systems genetic data to identify, visualize and prioritize functional assessment of key regulators remains a daunting task.

Here we present *CoffeeProt*, an easy to use online tool to enable the integrated analysis of proteomics data with pQTLs and molQTLs. *CoffeeProt* differs from existing tools by performing analyses primarily guided by the proteomics data, to discover protein-protein interactions to integrate with QTL data in interactive networks and SNP-protein association visualizations. Resources including the GWAS catalog and ENSEMBL variant effect annotations are integrated in the application, allowing for easy access to publicly available datasets and variant annotations. Ultimately, *CoffeeProt* enables the identification and prioritization of functionally relevant targets for follow up studies. *CoffeeProt* is accessible through an online user interface at www.coffeeprot.com, allowing analyses to be performed without programming knowledge and requires no software installation.

## MATERIALS AND METHODS

### CoffeeProt: an online tool for correlation and functional enrichment of proteome-wide systems genetics

*CoffeeProt* provides an easy to use and interactive interface allowing the seamless integration of proteomics data with QTL and molecular or phenotypic data. Correlation analyses are used to detect protein-protein associations which are further analyzed based on previously published interactions or shared annotations such as subcellular localization. The workflow is divided into proteomics processing, pQTL processing, molQTL processing, data analysis and plot export steps, as illustrated in **Figure 1**.

**Figure 1.**
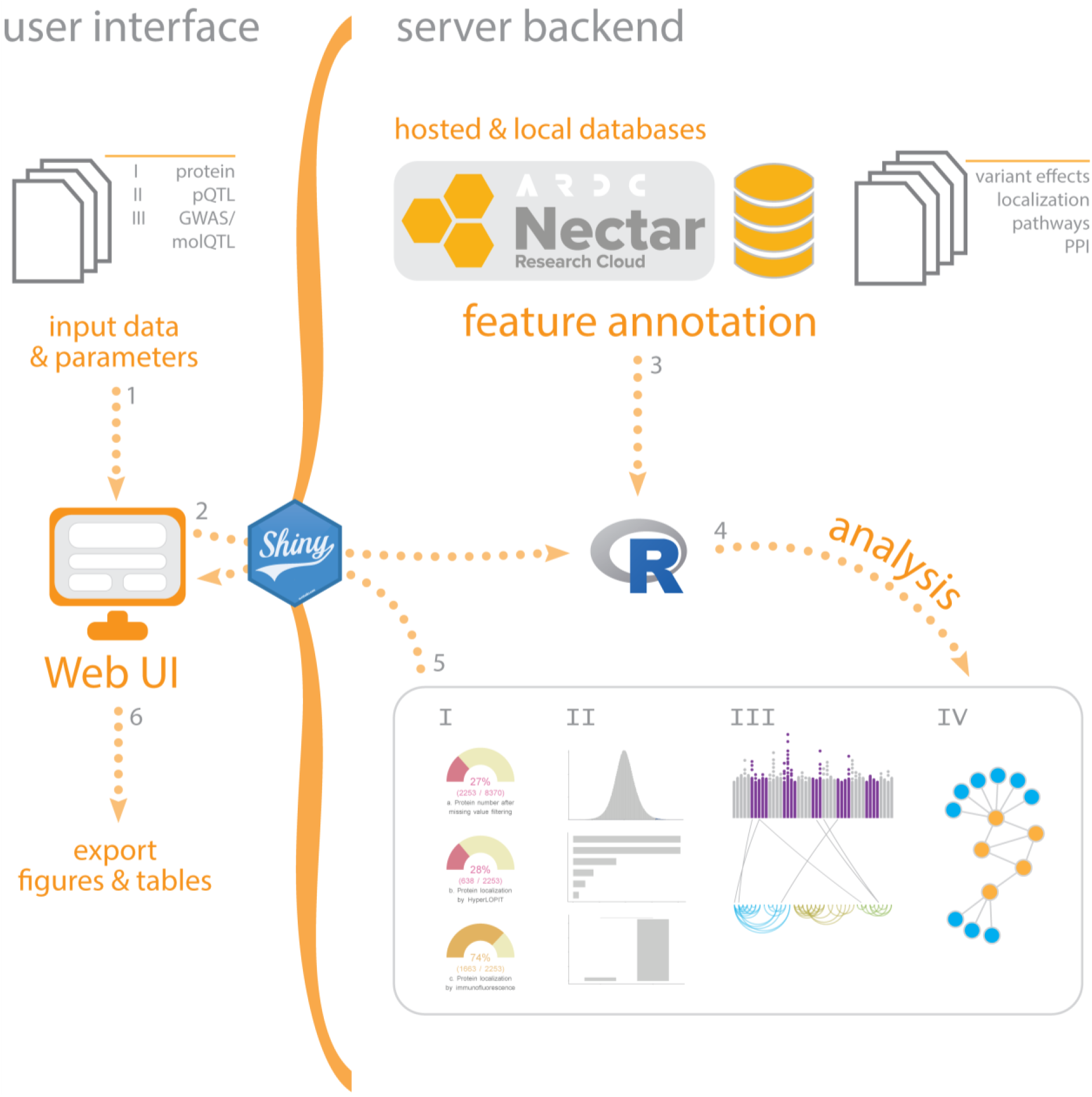
CoffeeProt workflow. The *CoffeeProt* workflow starts with users accessing the *CoffeeProt* web user interface at www.coffeeprot.com to upload datafiles and specify analysis parameters (1). The user interface and server backend running R are connected using the Shiny R package (2). Feature annotations are performed based on local databases included in *CoffeeProt* as well as remotely hosted databases on the Melbourne Research Cloud (3). User data is analyzed to perform summary statistics (I), correlation (II), interaction (III) and network (IV) analyses (4). The results are displayed in the web interface for result interpretation by the user (5). Finally, all individual tables and plots can be exported (6). (QTL: quantitative trait loci; PPI: protein-protein interaction).

### Data input & processing

The first step of the *CoffeeProt* workflow involves to user uploading input proteomics, pQTL and molQTL datasets using the web interface. The required format of these datasets is shown in **supplementary tables 1-3**. Example files are also available to download from the *CoffeeProt* welcome page. Datasets can be uploaded as comma separated .csv files, tab separated .txt files or Excel files in .xls or .xlsx formats. *CoffeeProt* automatically detects the file format and validates the dataset prior to data processing.

Proteomics data must be uploaded as a matrix in which the first column contains protein identifiers, and the remaining columns contain quantitative measurements. The accepted protein identifiers are gene names, Uniprot identifiers and ENSEMBL gene identifiers. *CoffeeProt* automatically detects the identifiers and converts them to gene names during the processing steps since these identifiers are required for several downstream analyses. Missing values in the proteomics data can be addressed by imputation or by removing proteins of which the number of missing values exceeds the cut-off defined by the user. Taylor *et al.* have analyzed the effects of missing values and imputation methods on the outcomes of correlation analyses in mass spectrometry-based -omics datasets (15). The authors showed that the percentage of false positive correlations increases whilst true positives decreases based on the level of missingness in the original dataset, irrespective of the imputation method used (15). Based on these observations the default missing value cut-off has been set to 20% in *CoffeeProt* but can be changed by the user. Following this filtering step, missing values are imputed using the MinDet function and default parameters from the imputeLCMD R package. Pair-wise protein-protein correlation or co-regulation analyses are performed using the Pearson correlation coefficient (PCC) or biweight midcorrelation (bicor) as implemented in the WGCNA R package (16). Following the correlation analysis and correction for multiple hypothesis testing with either Benjamini-Hochberg or Bonferroni, a list of protein pairs and their co-regulation metrics is created including the p-value, q-value and correlation coefficient.

For the pQTL data upload the matrix should contain separate columns containing (1) RefSeq identifiers (rsIDs), (2) SNP location, (3) SNP chromosome, (4) gene names, (5) gene start location, (6) gene end location, (7) gene chromosome, (8) a measure of significance and (9) a proxy or grouping column. Regarding the SNP and gene location columns, measures such as physical position (bp) or genetic distance (cM) are accepted. Non-cumulative measures (e.g. a measure that starts at zero for each chromosome) will be converted to a cumulative relative SNP location to allow the generation of Manhattan plots. The measure of significance column should contain either p-values or logarithm of the odds ratio (LOD) scores. Filtering of the pQTL dataset is allowed using a single measure of significance, or separate cut-offs for each QTL proxy. A proxy can be specified to allow separate cut-offs when the dataset contain both SNPs located near the gene it affects (local or *cis*-pQTLs) and SNPs that affect genes located in other regions of the genome (distant or *trans*-pQTLs). Due to the large file sizes common in QTL files, it is recommended to filter QTL data prior to uploading to *CoffeeProt*, to reduce the time required to upload data.

The third type of data upload is the association to phenotypic or molecular traits (GWAS or molQTLs). Data format is like the pQTL upload columns mentioned above. However, instead of (2) gene names, a molecule or phenotypic trait is listed as associated to the genetic variants. For the molQTL data upload the matrix should contain separate columns containing (1) RefSeq identifiers (rsIDs), (2) phenotypic trait(s), (3) SNP location, (4) SNP chromosome, (5) a measure of significance and (6) a grouping column. The use of molQTL/GWAS in *CoffeeProt* allows for the discovery of SNP-protein associations that share associations with molecular or clinical phenotypes. *CoffeeProt* enables easy access to publicly available data from the GWAS Catalog (17) for instances where the user only has access to proteomics and pQTL data. The molQTL tab in *CoffeeProt* contains a table listing all published studies in the GWAS Catalog, including the journal article title, publication date and traits investigated. A user can simply select a study of interest and click the download button to retrieve a list of variants associated to phenotypes that are directly usable as input molQTL data in *CoffeeProt*. The GWAS Catalog representational state transfer (REST) application programming interface (API) is accessed through the *gwasrapidd* R package (18). First all datasets related to the user selected study are retrieved using the get_associations function, followed by variant annotation using the get_variants function. Finally, the data table is trimmed to retain the six columns required for further analysis in *CoffeeProt.*

### Annotation

Imported proteins are annotated using subcellular localizations from the Cell Atlas as determined using immunofluorescence (19). All protein-protein correlation pairs are searched against the STRING database (20) as well as the CORUM (21) and BioPlex 3.0 (22) protein-protein interaction databases to detect previously reported associations. For the interactions in the BioPlex 3.0 database we only considered the top 4^th^ percentile (4727/118162) as correlated protein-protein interactions based on a previous publication by Huttlin *et al.* showing significant overlap between CORUM protein-protein interactions and the top 4^th^ percentile of the BioPlex interactions (23). A PostgreSQL database (v 11.3) hosted on the Nectar Research Cloud is used for the annotation of RefSeq identifiers in the pQTL datasets. This database contains both rsIDs and variant effects defined by the Sequence Ontology (24) which were retrieved from the latest ENSEMBL variation database (v100) (ftp://ftp.ensembl.org/pub/current_variation/vcf/) (25). Additionally, the rsIDs are assigned variant consequence impact ratings as used by variant annotation tools such as snpEff (26). Impact ratings refer to the disruptive effects the variant has on the functioning or effectiveness of a protein. High-impact variants are likely to cause protein truncation, loss of function or the triggering of nonsense mediated decay. A moderate-impact variant consequence is non-disruptive but may cause changes in protein activity or function such as an inframe insertion or deletion. A low-impact variant in unlikely to cause a change in protein behavior such as a synonymous variant. Finally, modifier-impact variant consequences affect non-coding areas are also annotated and these may influence the expression of the protein such as variants located in transcription factor binding sites or other regulatory elements (26).

### Analysis & visualization

*CoffeeProt* allows the customization of figures with user-selected cut-offs and the interactive selection of target proteins, complexes, and phenotypes or molecular traits. These data are integrated via network visualizations to ultimately understand how genetic variants are associated to protein complexes and different biological layers or phenotypes. *CoffeeProt* initially displays several plots related to the filtering and annotations of each of the datasets uploaded by the user. For proteomics datasets, multiple gauge charts are produced to highlight the number of proteins filtered by the missing value cut-off, and the number of proteins annotated by protein localization. Similarly, for QTL datasets multiple donut charts highlight the distribution of QTLs annotated by the proxy, grouping column, variant effects and variant impact ratings.

The correlation summary analysis tab contains visualizations related to the protein-protein correlation analysis. Histograms show the distribution of correlation coefficients or correlation q-values, highlighting the proportion that are associated based on user-defined cut-offs. Additionally, the number of correlated partners per protein is shown to highlight the proportion of highly connected proteins in the data set. To assess the quality of co-regulation data, enrichment analysis is performed in the Database analysis tab using the protein annotations of the protein-protein correlation pairs. Co-regulated proteins are expected to be enriched in the same cellular location and the same protein complexes as previously shown by Kustatscher *et al.* (27). The interactive nature of *CoffeeProt* allows users to adjust coefficient or q-value cut-offs to ensure functionally related protein pairs are significantly enriched.

*CoffeeProt* has several functionalities to explore the interactions between proteins and SNPs focused on the integration of several types of data. First, a Manhattan plot is created using the uploaded pQTL data, highlighting the loci associated with the abundances of specific proteins. Next, the protein complexes are organized to allow their visualization on a linear scale by sorting complexes by their number of subunit proteins and ordering the proteins within complexes by the number of correlations. These protein-protein correlations are shown as arc diagrams and are labelled by the complex name as it appears in the CORUM database. Alternatively, *CoffeeProt* allows the user to select a protein complex of interest, allowing a detailed look into its interactions and associated genetic variants. Finally, the connections between proteins and SNPs are annotated and colored to visually distinguish c*is*-QTLs and *trans*-QTLs or by variant effects and variant impact ratings.

Interactive network plots can be created using one or multiple biological layers to identify functionally relevant interactions. Networks can be created based on all measured proteins, all proteins with at least one QTL association or proteins with interactions previously reported in the CORUM or BioPlex 3.0 databases. The CORUM database already contains pre-defined networks, annotated with an identifier and description. For BioPlex 3.0, networks were produced from the top 4^th^ percentile of protein interactions, where all direct interactions with a target protein were considered to be part of that network. Protein-protein correlation analyses can lead to several million potential protein pairs, even in common proteomics datasets. Such an analysis could lead to 10,000-100,000 positively correlated protein pairs, depending on the criteria of the researcher. Filtering protein pairs to those affected by shared QTLs or genetic regions provides greater confidence that these associations are not spurious. pQTL data can be added to these network plots to highlight proteins affected by genetic variation. If molQTL or phenotypic data is present this can also be added. All QTLs can be shown as individual nodes for each rsID, or by combining rsIDs per chromosome. Summarizing QTLs by chromosome aids network visualization to collapse a group of SNPs e.g in linkage disequilibrium affecting a protein or trait. Bait-networks are produced by selecting a single protein or phenotype of interest for which all direct interactions are visualized. A significant feature of *CoffeeProt* is the identification and visualization of associations between proteins and phenotypes based on shared co-mapped SNP rsIDs. This is important because it identifies potential causative SNPs regulating both protein abundance and phenotypic or molecular traits for functional validation.

### Reporting

After completing the analysis workflow, it is possible to download all figures and tables produced by the user. The several datasets required to produce all images can be exported as .csv files for further analysis outside of *CoffeeProt*. All plots created in the tool can be directly exported in various formats, including vector-based images in .svg and .pdf formats and in various dimensions. The interactive network plots can be downloaded as .html files.

### Web server implementation

*CoffeeProt* was developed using the R programming language (28) for the back-end and relies on the *shiny* package for the web server front-end in addition to HTML, CSS and JavaScript. Several *tidyverse* packages are used for data wrangling and data visualization (29). Furthermore, *CoffeeProt* relies on the *WGCNA* package for the Pearson’s and bicor correlation analyses (8). Circos and interactive network plots are created using the *circlize* (30) and *networkD3* R packages respectively. The GWAS Catalog is accessed using the *gwasrapidd* package (18). *CoffeeProt* is deployed on the Nectar Research Cloud and Melbourne Research Cloud, utilizing hypervisors built on AMD EPYC 2 (base CPU clock speed 2.0Ghz, burst clock speed 3.35Ghz) and running Ubuntu 18.04.

## RESULTS

### CASE STUDY: An integrative systems genetic analysis of mammalian lipid metabolism

To illustrate the functionalities of the *CoffeeProt* tool, we analyzed data previously published by Parker *et al.* (14). In this study, mass spectrometry-based liver proteomic and lipidomic data were measured in the Hybrid Mouse Diversity Panel (HMDP) and integrated with genomic data via QTL analysis to identify genetic variants associated to proteins and lipids. For this case study we downloaded the supplemental information (https://www.nature.com/articles/s41586-019-0984-y#Sec34) and simply adapted the tables to be compatible with the *CoffeeProt* tool. The proteomics data were filtered to exclude proteins with more than 20% missing values amongst the samples, leaving a dataset containing 2,253 proteins in 306 mouse liver samples (**Figure 2A**). Among these proteins, 1,663 (74%) were annotated by protein localization as determined using immunofluorescent staining (**Figure 2B**).

**Figure 2.**
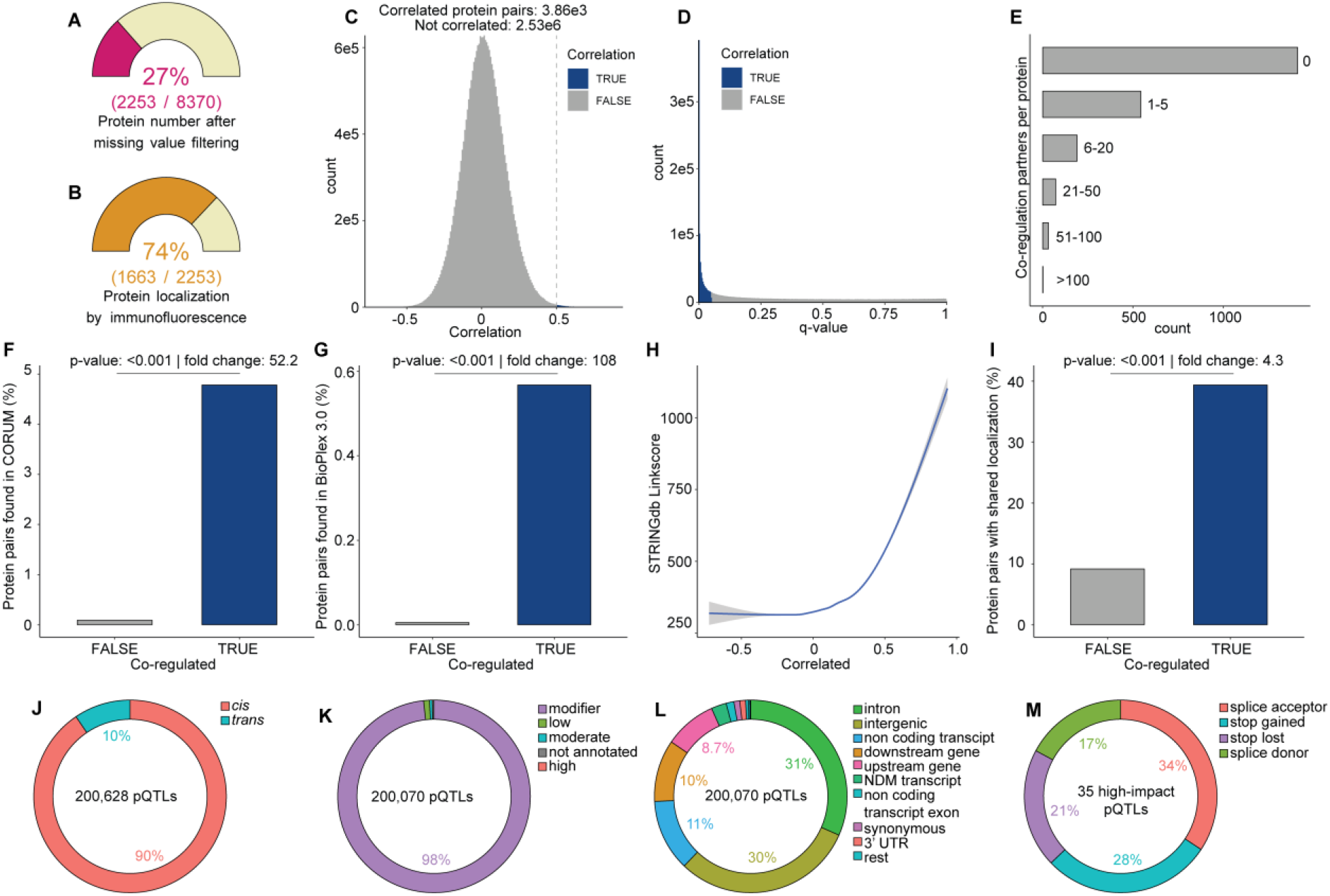
*CoffeeProt* pre-processing & correlation summary. (A) Percentage of proteins filtered by user-specified missing value cut-off. Percentage of proteins annotated with Cell Atlas protein localization as determined by (B) immunofluorescent staining. The number of co-regulated protein pairs after filtering for user-specified correlation (C) or q-value (D) cut-offs. The number of protein pairs that meets the criteria is displayed in both plots. The number of co-regulation partners per proteins (E) has been determined based on the user-specified correlation cut-off. The data is binned to group proteins with 0, 1-5, 6-20, 21-50, 51-100 and >100 partners. Enrichment of previously reported protein-protein interactions as reported in the STRING (F), CORUM (G) and Bioplex 3.0 (H) databases. Enrichment of protein pair co-localization (I) was determined based on Cell Atlas localization determined by immunofluorescent staining. A Chi-squared test was performed to determine the significance of enrichments. pQTL annotation distributions by proxy (J), variant impact (K) and variant effect (L, M).

#### Protein Correlation Summary

We performed a protein-protein correlation analysis to obtain biweight midcorrelation (bicor) coefficients, followed by the Benjamini-Hochberg procedure to obtain adjusted p-values (q-values). Even a relatively small proteomics dataset will lead to several million unique protein pairs, indicating the necessity to differentiate between truly associated protein pairs and sporadic correlations. Here we considered proteins with a bicor coefficient greater than 0.5 and q-value < 0.05 to be correlated. This correlation coefficient cut-off led to 0.15% (3.86e3 out of 2.53e6) of the protein-pairs to be defined as correlated (**Figure 2C-D**). Using these cut-offs, most proteins have no or a small number (1-5) of co-regulation partners (**Figure 2E**). A smaller group of proteins has many co-regulation partners, indicating proteins that are likely central in one or multiple protein complexes.

#### Protein Correlation Database enrichment

Analyzing the fraction of previously reported protein pairs in the correlation matrix based on the CORUM and BioPlex 3.0 databases revealed an enrichment of database membership by 52.2- and 108-fold, respectively compared to the non-correlated pairs (**Figure 2F-G**). Similarly, the higher positive correlation coefficients were associated with increased STRINGdb linkscores (**Figure 2H**). These results indicate that highly correlated protein pairs discovered in this workflow are genuine associations, based on previously reported protein-protein interactions. Consistent with previous co-regulation network analysis, top CORUM protein complexes identified included mitochondrial complex I, ribosome, proteasome, and spliceosome. Furthermore, it is expected that co-regulated proteins have a larger proportion of overlapping annotations. Indeed, among the correlated protein pairs almost 40% share a subcellular localization compared to less than 10% for the non-correlated pairs (**Figure 2I**).

#### pQTL and lipid-QTL summary

A total of 200,628 SNPs were associated to the abundance of 978 proteins of which 90% were associated to 789 proteins via *cis*-pQTLs (+/− 10 mb, p < 1e-4) and 10% were associated to 436 proteins via *trans*-pQTLs (p < 1e-7) (**Figure 2J**). Furthermore, 1,336 SNPs were identified within 336 genes and were also associated to the expression of these corresponding proteins via *cis*-pQTLs. Although most variants are modifier-impact (>98%), small numbers of low- (1982, 1%), moderate- (913, 0.4%), and high-impact (35, 0.02%) SNPs were found (**Figure 2K**). Annotating these SNPs with ENSEMBL variant effects (**Figure 2L**) revealed that most SNPs are intron (31%) or intergenic (30%) variants. A total of 35 high-impact SNPs were associated to protein abundance via variants in stop codons or splicing sites (**Figure 2M**).

We also performed variant effect analysis on 1,808 SNPs that were associated to 89 lipid species and 17 lipid classes. The dataset contained 1338 associations with lipid classes and 6694 associations with individual liver lipids.

#### Integration of co-regulation networks, pQTLs and lQTLs

To visualize the associations between genetic variation and co-regulated protein networks, *CoffeeProt* produces QTL-protein plots. A Manhattan plot is created using pQTL data and loci are linked to co-regulated networks that are depicted as arc diagrams. The lines linking the Manhattan plot and arc diagrams can be colored with either the *cis-/trans*-proxies, variant effects or impact ratings. The Manhattan plots can be filtered for specific chromosomes and/or co-regulated complexes can also be filtered for CORUM or BioPlex protein-protein interaction complexes. **Figure 3A** displays genome-wide variants associated with co-regulated CORUM complexes annotated with *cis-/trans*-proxies. **Figure 3B**, Mitochondrial Complex I (CORUM ID 382) has been selected to investigate variants associated to precise subunits which have now been annotated with impact ratings. In this example, a high-impact *trans*-pQTL associated to NDUFS2 has been identified and analysis of the pQTL summary page identifies a trans-acting variant in a predicted splicing acceptor site (rs27441698) within the *Sptbn5* gene.

**Figure 3.**
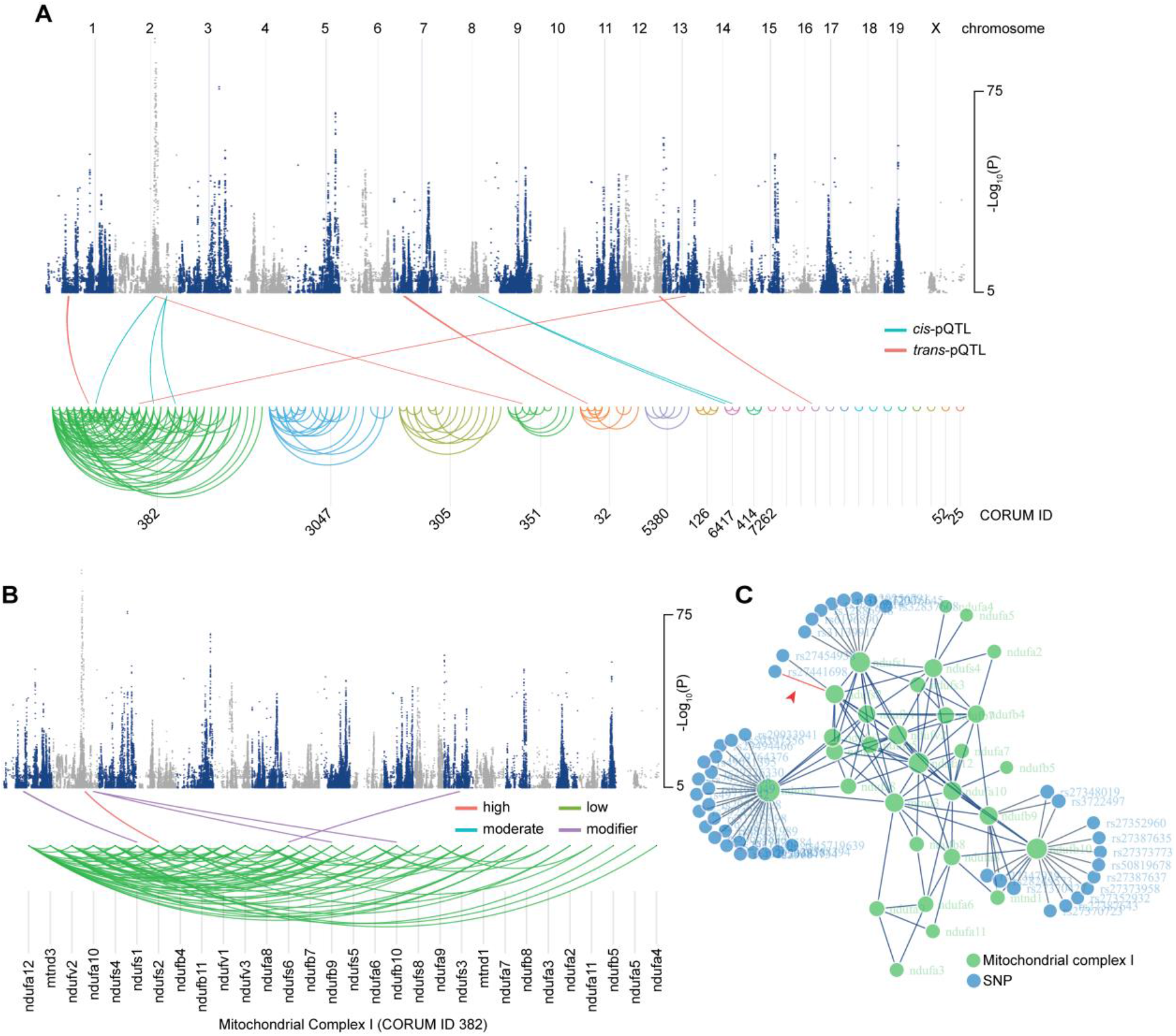
QTL-protein associations of mitochondrial complex I. (A) QTL-protein associations of all correlated CORUM protein networks or (B) only highlighting mitochondrial complex I. SNP-protein associations are colored by QTL type or SNP variant impact. (C) A network plot reveals the number of SNP associations for the proteins in the CORUM network.

We next demonstrate *CoffeeProt’s* ability to generate multi-omic visualizations via interactive networks. Users can select various co-regulated networks based on CORUM or BioPlex and integrate associated genetic variants as nodes with edges annotated by proxy, variant effects or impact ratings. **Figure 3C** displays an alternative network view of the pQTLs associated to Mitochondrial Complex I with the example high-impact association mentioned above shown in red and indicated by a red arrowhead. In addition to a protein co-regulation-centric network analysis, users can choose to focus on a specific protein and perform a bait network analysis that includes associations to molecular or phenotypic traits. **Figure 4A** highlights genetic variants associated to ACOT13, an important enzyme regulating hepatic lipid metabolism via the hydrolysis of fatty acyl-CoAs (31). *CoffeeProt* identified several co-mapping *cis-* pQTLs to ACOT13 and lQTLs to long chain phosphatidylcholine (PC), phosphatidylinositol (PI) and phosphatidylethanolamine (PE) lipid species. Furthermore, intragenic SNPs were identified including several in the first and second introns of *Acot13* that were associated to both ACOT13 and lipid abundance supporting previous roles of this protein in lipid metabolism. *CoffeeProt* also allows users to perform a molecular or phenotypic-centric network analysis to investigate potential upstream protein and genetic regulators. **Figure 4B** highlights co-mapping pQTLs and lQTLs to hepatic cholesterol ester abundance. Two previously characterized regulators of cholesterol metabolism are highlighted with red arrows including CYP51, a monooxygenase catalyzing the first step in the conversion of lanosterol into cholesterol (32), and TMEM97, a lysosomal protein associated with the regulation of low-density lipoproteins (LDLs) and cholesterol ester metabolism (33). These examples demonstrate *CoffeeProt’s* ability to identify previously reported regulators of lipid metabolism and highlight many more uncharacterized associations for future functional validation.

**Figure 4.**
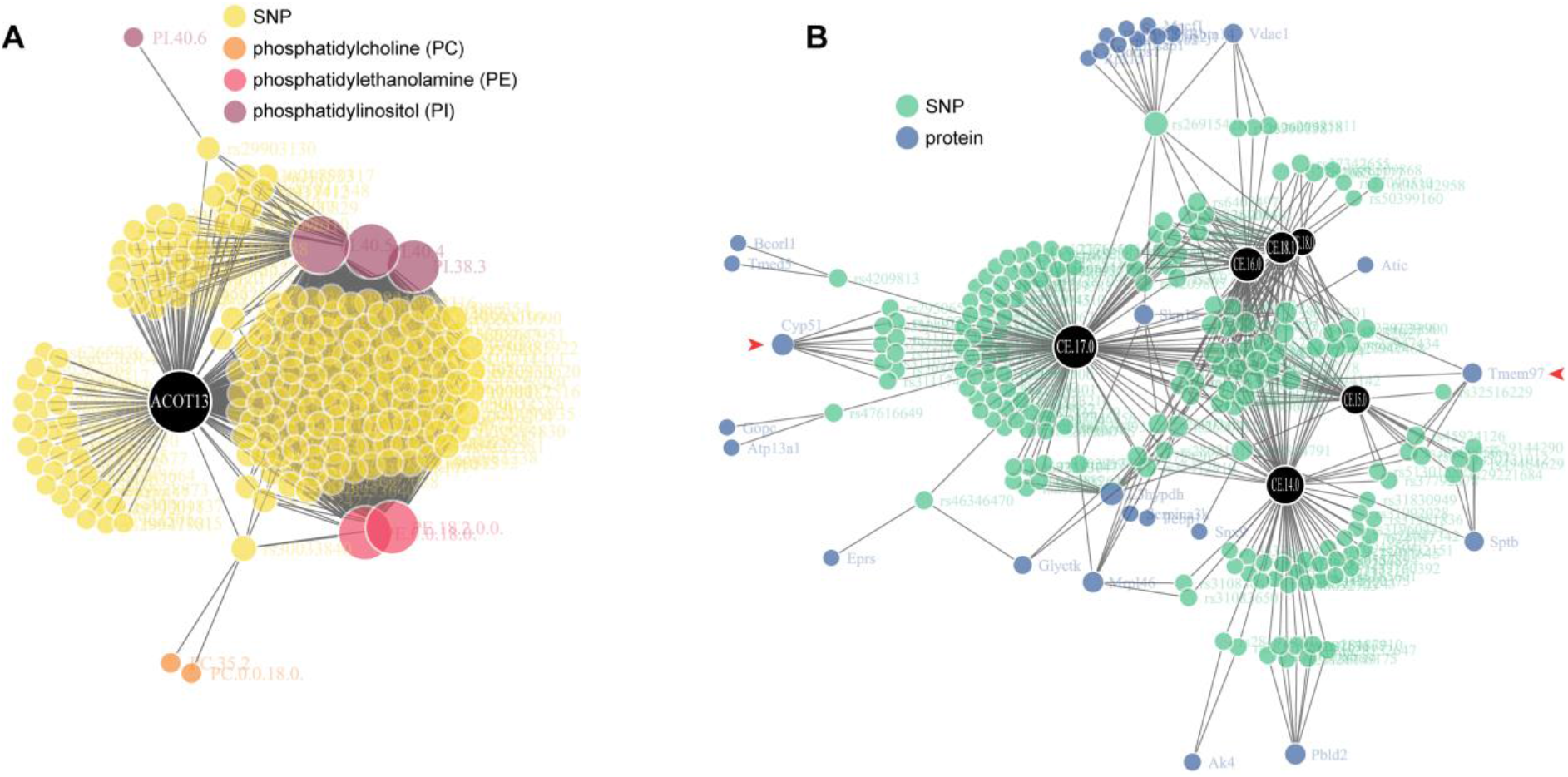
Regulators of plasma lipids. (A) Protein bait-network shows all phenotypes associated with ACOT13 by highlighting shared SNPs. (B) Phenotype bait-network in which CE 14:0. CE 15:0. CE 16:0, CE 17:0. CE 18:0 and CE 18:1 were selected as targets. (CE: cholesterol ester, PC: phosphatidylcholine, PE: phosphatidylethanolamine, PI: phosphatidylinositol)

### CASE STUDY: Genomic atlas of the human plasma proteome

As a second case study we used *CoffeeProt* to analyze genetic associations of the human plasma proteome from the INTERVAL study published by Sun *et al*. (34). In this study, SOMAscan-based proteomics was used to quantify over 3,200 proteins in the plasma of 3,301 healthy participants and integrated with genomic data via pQTL anlaysis. Access to the proteomics and pQTL data was provided upon request through the European Genotype Archive (accession number EGAS00001002555). The data was simply pre-processed to filter columns and uploaded directly to *CoffeeProt* for further analysis. The proteomics data was filtered to exclude proteins with more than 20% missing values amongst the samples, leaving a dataset containing 3,257 proteins (**Figure 5A**). Among these proteins, 1,863 (57%) were annotated by protein localization as determined using immunofluorescent staining as reported in the Human Protein Atlas (19) (**Figure 5B**).

**Figure 5.**
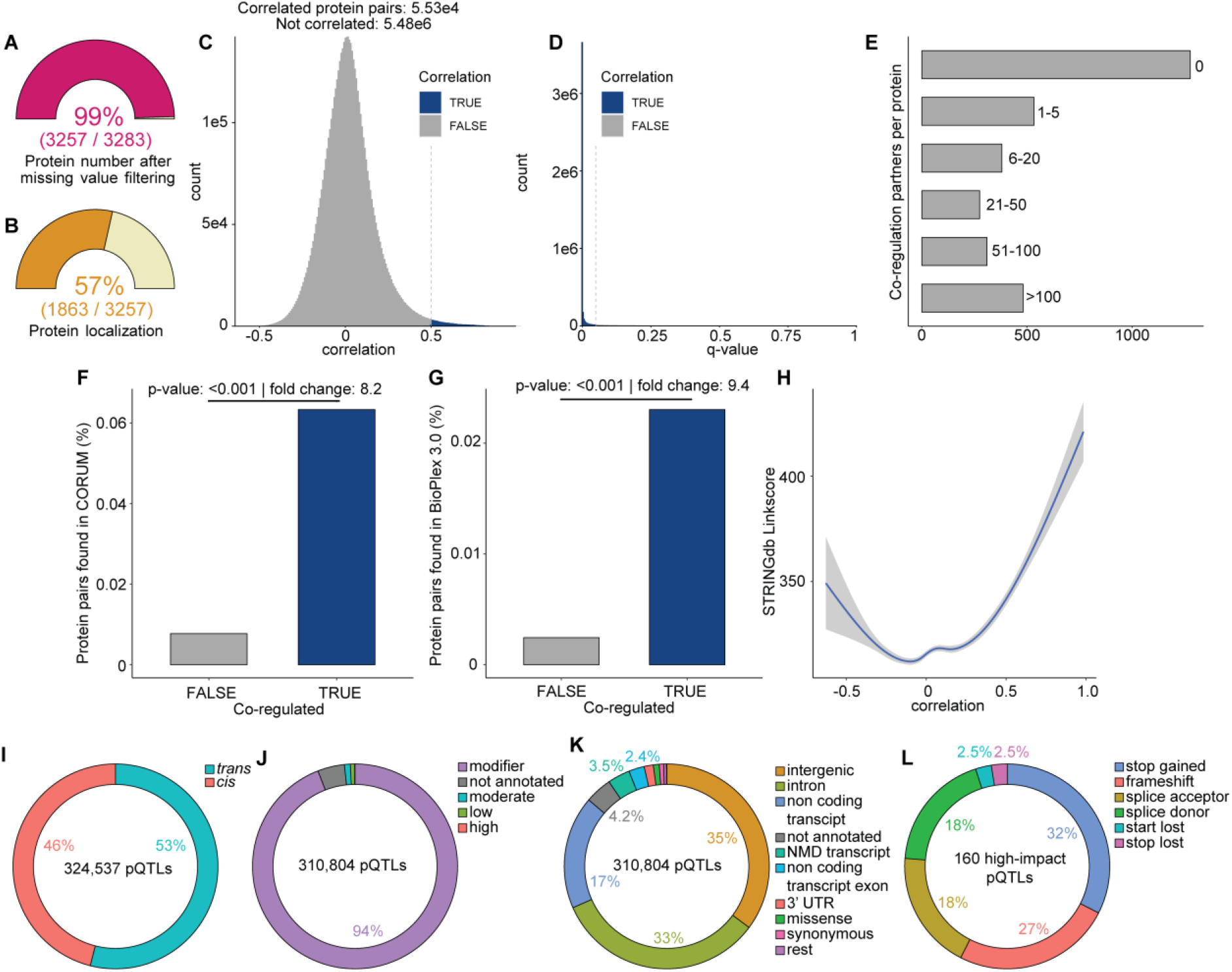
*CoffeeProt* pre-processing & correlation summary. (A) Percentage of proteins filtered by user-specified missing value cut-off. Percentage of proteins annotated with Cell Atlas protein localization as determined by (B) immunofluorescent staining. The number of co-regulated protein pairs after filtering for user-specified correlation (C) or q-value (D) cut-offs. The number of protein pairs that meets the criteria is displayed in both plots. The number of co-regulation partners per proteins (E) has been determined based on the user-specified correlation cut-off. The data is binned to group proteins with 0, 1-5, 6-20, 21-50, 51-100 and >100 partners. Enrichment of previously reported protein-protein interactions as reported in the STRING (F), CORUM (G) and Bioplex 3.0 (H) databases. A Chi-squared test was performed to determine the significance of enrichments. pQTL and molQTL annotation distributions by proxy (I), variant impact (J) and variant effect (K, L).

#### Protein Correlation Summary

Like the first case study, we performed protein-protein correlation using the biweight midcorrelation (bicor), followed by the Benjamini-Hochberg procedure. We considered proteins with a bicor coefficient > 0.5 and q-value < 0.05 to be correlated. This correlation coefficient cut-off led to 1% (5.53e4 out of 5.48e6) of the protein-pairs to be defined as correlated (**Figure 5C-D**). Using these cut-offs, most proteins have no or a small number (1-5) of co-regulation partners (**Figure 5E**). A smaller group of proteins have many co-regulation partners, indicating proteins that are likely central in one or multiple protein complexes.

#### Protein Correlation Database enrichment

Analyzing the fraction of previously reported protein pairs in the correlation matrix based on the CORUM and BioPlex 3.0 databases revealed an enrichment of database membership by 8.2- and 9.4-fold, respectively compared to the non-correlated pairs (**Figure 5F-G**). However, it should be noted that only a small fraction of the quantified plasma proteome was mapped to these databases presumably because protein-protein interaction analysis has previously focused on intracellular proteins. Similarly, the higher positive correlation coefficients were associated with increased STRINGdb linkscores (**Figure 5H**). These results indicate that despite the low annotation of plasma protein complexes, highly correlated protein pairs discovered in this workflow are genuine associations, based on previously reported protein-protein interactions.

#### pQTL summary

The pQTL data as processed by Sun *et al.* were filtered by p-value (p < 1e-4) and all resulting tables were merged, resulting in a table of ~5,000,000 pQTLs. The SOMAmer identifiers in the QTL data were annotated by mapping to their respective gene names and locations (chromosome and bp) on the GRCh37/hg19 reference genome. UniProt identifiers were converted to gene names using the AnnotationDbi and org.Hs.eg.db R packages. GENCODE human release 19 (CRCh37.p13) was used to retrieve genomic locations for all measured proteins. We defined SNPs located < 1MB from the start of the target gene as *cis*-pQTLs, whereas all other SNPs are defined as *trans*-pQTLs. The resulting pQTL file was uploaded to *CoffeeProt* for further filtering, analysis and visualization. Both *cis-* and *trans-* pQTLs were filtered to p < 1e-11 consistent with the original publication resulting in the identification of 324,537 SNPs associated to the abundance of 1,409 proteins of which 46% were associated to 530 proteins via *cis*-pQTLs and 53% were associated to 1,066 proteins via *trans*-pQTLs (**Figure 5I**). Although most variants were modifier-impact (94%), small numbers of low- (2215, 0.6%), moderate- (3027, 0.9%), and high-impact (160, 0.008%) SNPs were found (**Figure 5J**). The majority of SNPs annotated with the ENSEMBL variant effects were intergenic (35%), intron (33%), or non-coding transcript variants (17%) (**Figure 5K**). The high-impact variants were categorized as stop gained- (52), frameshift- (40), splice donor- (30), splice acceptor- (30), start lost- (4) or stop lost (4) variants (**Figure 5L**).

#### Integration of co-regulation networks, pQTLs and GWAS

We initially investigated intragenic SNPs that were associated to the abundance of co-regulated protein networks. **Figure 6A** displays a QTL-Protein plot with *cis-*acting pQTLs associated to co-regulated protease complement factors CFB, CFH, CFI and APCS. *CoffeeProt* identified previously characterized high-impact SNPs including rs4151667 and rs641153, resulting in non-synonymous L9H and R32Q variants of CFB, respectively (35,36). Several co-mapping *cis-*acting variants in the first intron of CFH were also associated to the abundance of CFB via trans-pQTLs. Given the abundance of these two proteins are correlated, these data suggest genetic regulation of CFH subsequently regulates the abundance of CFB. Hence, *CoffeeProt* allows the prioritization of further causal analysis such as downstream mediation analysis. Direct integration of data from the GWAS Catalog revealed co-mapping SNPs associated to several clinical phenotypes such as a previously characterized role of these variants in macular degeneration (**Figure 6B**). *CoffeeProt* also enabled the discovery of several uncharacterized co-regulated networks associated to clinical phenotypes. For example, intragenic *cis*-pQTLs were associated to the abundance of Tenascin (TNC) and co-mapped to *trans-*pQTLs in a c-oregulated network of EVA1C, FAM163A and SFTPC (**Figure 6C**), The presence of these shared SNPs suggests that some of these proteins are affected by SNPs related to another gene or chromosome. Indeed, EVA1C and FAM163A are not affected by SNPs located on their respective chromosomes (chr21 & chr1) but only by SNPs located on chromosomes 8 & 9 at the genomic locations of TNC and SFTPC (**Figure 6D**). On the other hand, the SNPs near TNC and SFTPC are associated with both TNC and SFTPC. These findings suggest that variations in TNC and SFTPC affect EVA1C and FAM163A, but not vice versa. Both TNC and FAM163A, which show a significant protein-protein interaction (bicor coefficient 0.84, q-value < 0.001), and show a common association with body mass index through SNP rs12376870. Although the link between TNC and BMI was previously known (37), the functions of FAM163A are not well-defined. However, recent experimental data in C57BL/6N mice lacking *Fam163a* from the International Mouse Phenotyping Consortium (38) has shown significant associations with decreased total body fat amount, decreased body length, decreased circulating glucose levels and increased lean body mass. These examples demonstrate CoffeeProt’s ability to rapidly integrate and visualize co-regulated network analysis, pQTL and GWAS data to aid functional interpretation and prioritize candidate validation.

**Figure 6.**
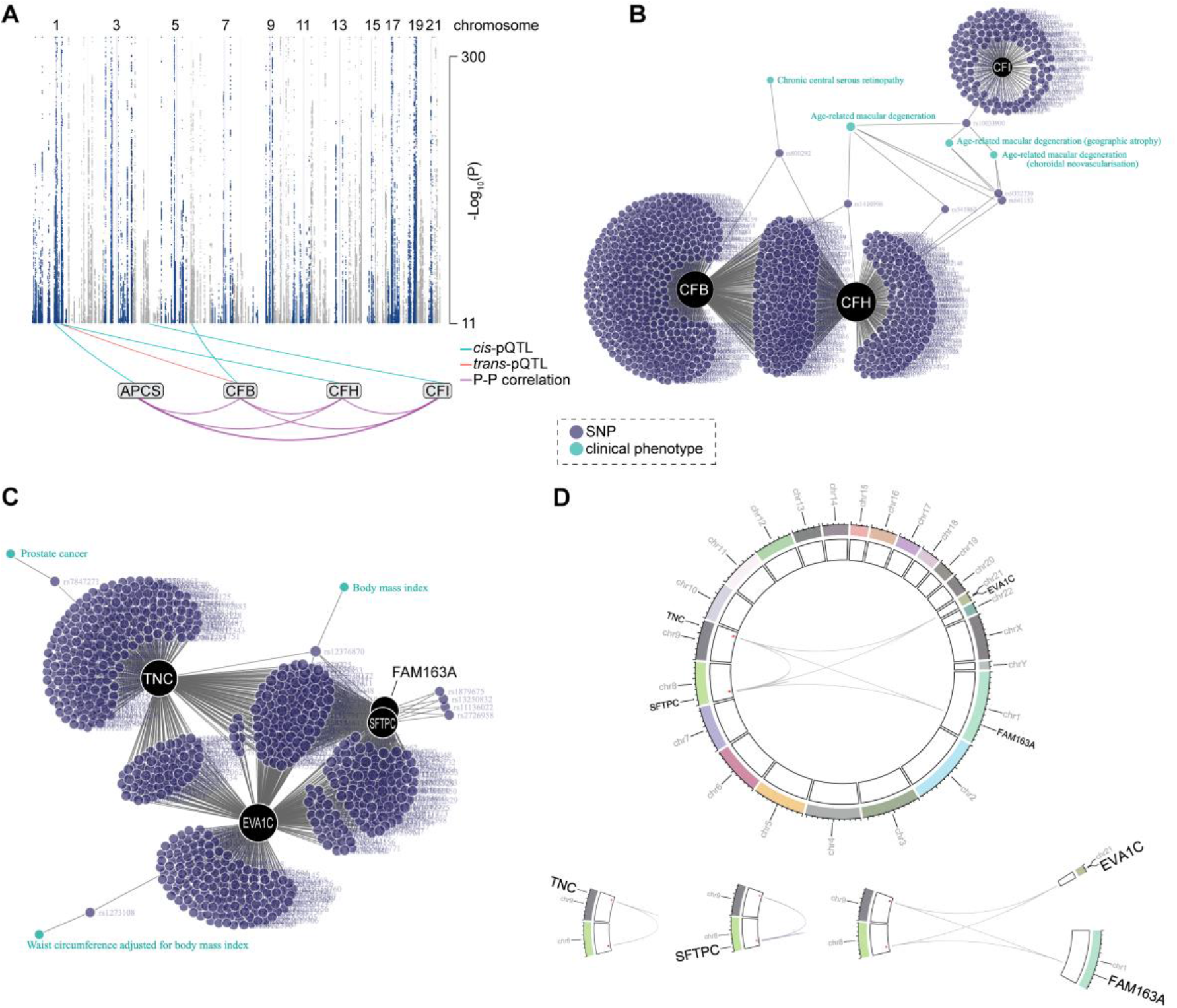
Integration of co-regulation networks, pQTL and GWAS in human plasma samples. (A) A SNP-protein association plot highlights *cis-* and *trans-*pQTLs associated with interacting proteins. (B) The CFB-CFH-CFI network reveals shared SNPs between the proteins and clinical phenotypes from the GWAS catalog. (C) Mapping the INTERVAL proteomics data to previously published phenotypes from the same study produced a network of TNC, EVA1C, SFTPC and FAM163A, including their associated SNPs and phenotypes. (D) The Circos plot shows the genomic locations of the SNPs affecting the proteins TNC, EVA1C, SFTPC and FAM163A. The dots in the second ring indicate the locations of the SNPs with the links in the center connecting the SNPs to the affected genes / proteins.

We next used *CoffeeProt* to perform a clinical phenotypic-centric network analysis by integrating previous GWAS analysis to plasma pQTLs. First, we investigated genetic variants associated to 36 blood cell traits that were directly downloaded from the GWAS Catalog in CoffeeProt (39) and identified co-mapping plasma pQTLs. **Figure 7A** displays associations to various blood cell counts and other clinical measures including mean corpuscular volume (MCV) and hemoglobin content. This interactive network provides an exciting glimpse into the complex genetic regulation of human blood and plasma proteome revealing several known and novel associations. Red blood cell associations are clustered on the left while other blood cells (granulocyte, leukocyte, basophil, eosinophil) are clustered on the right. The grouping of these cell types highlights the distinct groups of SNPs affecting the different blood cell types, but also SNPs shared between red and white blood cells. The subnetwork of red blood cell-related phenotypes (**Figure 7B**) shows considerable variant overlap between these traits, as well as large numbers of SNPs unique to each trait. **Figure 7C** highlights previously characterized *cis*-acting SNPs in the ABO locus including rs149037075 and rs10901252 which CoffeeProt has annotated as 3 prime UTR variants, and rs8176643, annotated as an intron variant. The variant rs8176643 was validated in a meta-analysis of SNPs in the ABO locus and their associations with red blood cell traits (40). Sun *et al.* previously identified ABO as one of the highly pleiotropic loci associated with more than 20 *trans*-associations (34). This agrees with a study by Emilsson *et al*, who. identified over 40 proteins via *trans*-pQTLs associated with variants located on the *ABO* gene, which were also associated with cardiovascular disease and hemostasis (41). The traits “hemoglobin measurement” and “erythrocyte count” are both associated with two separate SNPs (rs66782572 & rs3617) which are *cis*-pQTLs of the protein ITIH1 (**Figure 7C**). This protein is a known marker for high-altitude adaptation although its mechanism of action was not revealed (42). Furthermore, both SNPs are associated *in trans* to the proteins SIGIRR, JAKMIP3 and NAT1 in addition to the association of rs3617 with C15orf48 (NMES1). Indeed, searching the GWAS catalog reveals that the strongest associations of SNPs mapped to the ITIH1 gene, reported by Emilsson *et al.* (study ID GCST006585) (43), are blood protein concentrations of SIGIRR (1e^-300^), ITIH1 (5e^-160^), RBL2 (2e^-42^), ITIH3 (7e^-39^), NMES1 (C15orf48) (3e^-29^) and JAKMIP3 (2e^-25^). The SNP rs3617 is a missense variant of ITIH3 which could explain its *cis*-associations with ITIH1. The link between NMES1 (C15orf48), ITIH1 and erythrocyte counts is interesting since the functions of NMES1 in relation to blood cells are unknown as it is considered to be a marker for normal esophageal tissue (44). An independent proteomics study revealed altered concentrations of both ITIH1 and NMES1 as a response to differing oxygen concentrations, affecting human macrophage polarization and functioning (45). NMES1 was one of several markers associated with DC1 and DC17 effector dendritic cells (46). Furthermore, it was shown that the transcript for NMES1 is a functional microRNA (miR-147) (47), and could be a potential transcription regulator of PDPK1 and subsequently AKT (48). miR-147 was also correlated with hemagglutinin mRNA expression, which is a glycoprotein involved in red blood cell agglutination and coagulation (49). Interestingly, the protein-protein correlation data showed no evidence of a direct interaction between ITIH1 and NMES1. This could be due to the regulation of *NMES1* by SNPs from various regions of the genome (Chr 3, Chr 19), rather than an exclusive regulation by the SNPs on *ITIH3* (data not shown). These SNPs on chromosome 19 regulating *NMES1 in trans* included rs10418046 and rs62143206, an upstream intergenic variant and an intron variant mapped to *NLRP12.* Taken together, these examples highlight the power of *CoffeeProt* to rapidly integrate, visualize and interrogate multiple associations to complex human traits.

**Figure 7.**
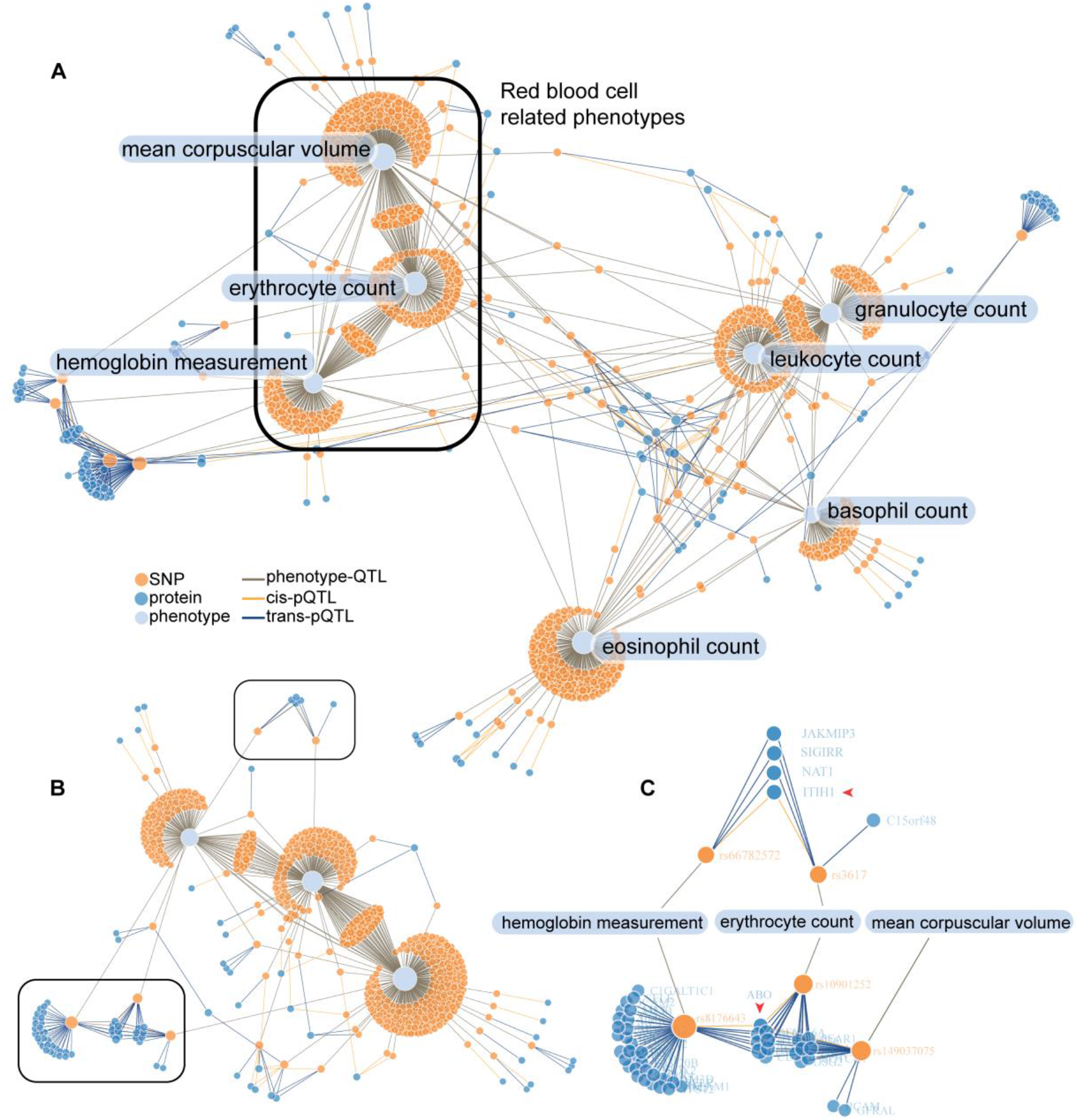
Regulators of blood measurements and cell counts. (A) Phenotype bait-network of blood measurements and cell counts from the INTERVAL cohort. The outlined subnetwork represents red blood cell-related phenotypes, which is highlighted in (B). The outlined boxes contain at least one protein in *cis*-pQTL associations with two or more SNPs. (C) Proteins associated with SNPs located on or near ABO and ITIH1. The red arrowheads indicate proteins with *cis*-pQTL associations. Link colors indicate the QTL types: phenotype-QTL (green), *cis*-pQTL (yellow), *trans*-pQTL (blue).

## DISCUSSION

We present *CoffeeProt*, a novel online tool for the correlation and functional enrichment of proteome-wide systems genetics. *CoffeeProt* is flexible and accepts proteomics data from various sources including mass spectrometry- and aptamer-based experiments. Users could complete the full *CoffeeProt* workflow within several minutes and without the need for any programming knowledge. The use of a dedicated database for SNP variant effect has allowed the rapid annotation of QTLs, allowing easy prioritization of associations based on predicted variant impacts. By allowing integrative analyses to be performed on proteomic-generated co-regulated network analysis, pQTL, molQTL, and phenotypic data, users can visualize and discover interactions and associations which may otherwise have been missed. Furthermore, mapping *cis-* and *trans-*pQTL data onto protein-protein co-regulated networks offers several advantages to investigate the genetic effects on protein complexes and potentially limit false positive associations by observing co-mapping SNPs. As shown in two case studies, *CoffeeProt* allows for the discovery of biologically relevant associations and genetic regulations. We highlighted regulators of lipid metabolism associated with phospholipid (ACOT13) and cholesterol ester concentrations (CYP51, TMEM97) in agreement with previous publications. Furthermore, using data from the INTERVAL study highlighted well-known (ABO) and uncharacterized (NMES1) associations with blood traits.

In summary, *CoffeeProt* is a valuable tool in proteome wide association studies introducing a variety of analyses and visualizations for the discovery of biologically relevant interactions and associations. We aim to support and continually improve *CoffeeProt* over the coming years based on user feedback and the changing bioinformatics requirements in the systems genetics field.

## DATA AVAILABILITY

*CoffeeProt* is accessible as a website (www.coffeeprot.com). The Parker HMDP data files used as case study 1 are included as demo files available to download through the *CoffeeProt* website. The data used in case study 2 (Sun *et al*.) are available through the European Genotype Archive (accession number EGAS00001002555). RefSeq identifiers and variant effects defined by the Sequence Ontology (24) which were retrieved from the latest ENSEMBL variation database (v100) (ftp://ftp.ensembl.org/pub/current_variation/vcf/). Subcellular protein localization information was downloaded from the Human Protein Atlas (19) (https://www.proteinatlas.org/download/subcellular_location.tsv.zip). GENCODE human release 19 (CRCh37.p13) was used to retrieve genomic locations for the pQTL data in the second case study. The CORUM database was downloaded from http://mips.helmholtz-muenchen.de/corum/#download. BioPlex 3.0 was downloaded from https://bioplex.hms.harvard.edu/interactions.php.

## SUPPLEMENTARY DATA

Supplementary Tables S1-3

## ACKNOWLEDGEMENT

This research was supported by use of the Nectar Research Cloud and by the University of Melbourne Research Platform Services. The Nectar Research Cloud is a collaborative Australian research platform supported by the National Collaborative Research Infrastructure Strategy. This work was funded by an Australian National Health and Medical Research Council Ideas Grant (APP1184363) and The University of Melbourne Driving Research Momentum program. We would like to thank the Melbourne Mass Spectrometry and Proteomics Facility of The Bio21 Molecular Science and Biotechnology Institute at The University of Melbourne for the support of mass spectrometry analysis. We would like to thank Adam S. Butterworth for providing access to the proteomics and pQTL data from the INTERVAL study. The contents of the published material are solely the responsibility of the individual authors and do not reflect the view of funding bodies.

## REFERENCES

1. Claussnitzer, M., Cho, J.H., Collins, R., Cox, N.J., Dermitzakis, E.T., Hurles, M.E., Kathiresan, S., Kenny, E.E., Lindgren, C.M., MacArthur, D.G. et al. (2020) A brief history of human disease genetics. Nature, 577, 179–189.

2. Civelek, M. and Lusis, A.J. (2014) Systems genetics approaches to understand complex traits. Nat Rev Genet, 15, 34–48.

3. Williams, E.G. and Auwerx, J. (2015) The Convergence of Systems and Reductionist Approaches in Complex Trait Analysis. Cell, 162, 23–32.

4. Ye, Y., Zhang, Z., Liu, Y., Diao, L. and Han, L. (2020) A Multi-Omics Perspective of Quantitative Trait Loci in Precision Medicine. Trends Genet, 36, 318–336.

5. Stacey, D., Fauman, E.B., Ziemek, D., Sun, B.B., Harshfield, E.L., Wood, A.M., Butterworth, A.S., Suhre, K. and Paul, D.S. (2019) ProGeM: a framework for the prioritization of candidate causal genes at molecular quantitative trait loci. Nucleic Acids Res, 47, e3.

6. Seldin, M., Yang, X. and Lusis, A.J. (2019) Systems genetics applications in metabolism research. Nat Metab, 1, 1038–1050.

7. Arneson, D., Bhattacharya, A., Shu, L., Makinen, V.P. and Yang, X. (2016) Mergeomics: a web server for identifying pathological pathways, networks, and key regulators via multidimensional data integration. BMC Genomics, 17, 722.

8. Langfelder, P. and Horvath, S. (2008) WGCNA: an R package for weighted correlation network analysis. BMC Bioinformatics, 9, 559.

9. Song, W.M. and Zhang, B. (2015) Multiscale Embedded Gene Co-expression Network Analysis. PLoS Comput Biol, 11, e1004574.

10. Argelaguet, R., Velten, B., Arnol, D., Dietrich, S., Zenz, T., Marioni, J.C., Buettner, F., Huber, W. and Stegle, O. (2018) Multi-Omics Factor Analysis-a framework for unsupervised integration of multi-omics data sets. Mol Syst Biol, 14, e8124.

11. Margolin, A.A., Nemenman, I., Basso, K., Wiggins, C., Stolovitzky, G., Dalla Favera, R. and Califano, A. (2006) ARACNE: an algorithm for the reconstruction of gene regulatory networks in a mammalian cellular context. BMC Bioinformatics, 7 Suppl 1, S7.

12. Chick, J.M., Munger, S.C., Simecek, P., Huttlin, E.L., Choi, K., Gatti, D.M., Raghupathy, N., Svenson, K.L., Churchill, G.A. and Gygi, S.P. (2016) Defining the consequences of genetic variation on a proteome-wide scale. Nature, 534, 500–505.

13. Keele, G.R., Zhang, T., Pham, D.T., Vincent, M., Bell, T.A., Hock, P., Shaw, G.D., Munger, S.C., de Villena, F.P.M., Ferris, M.T. et al. (2020) Regulation of protein abundance in genetically diverse mouse populations. bioRxiv, September 18, 2020, https://doi.org/10.1101/2020.1109.1118.296657.

14. Parker, B.L., Calkin, A.C., Seldin, M.M., Keating, M.F., Tarling, E.J., Yang, P., Moody, S.C., Liu, Y., Zerenturk, E.J., Needham, E.J. et al. (2019) An integrative systems genetic analysis of mammalian lipid metabolism. Nature, 567, 187–193.

15. Taylor, S.L., Ruhaak, L.R., Kelly, K., Weiss, R.H. and Kim, K. (2017) Effects of imputation on correlation: implications for analysis of mass spectrometry data from multiple biological matrices. Brief Bioinform, 18, 312–320.

16. Langfelder, P. and Horvath, S. (2012) Fast R Functions for Robust Correlations and Hierarchical Clustering. J Stat Softw, 46.

17. Buniello, A., MacArthur, J.A.L., Cerezo, M., Harris, L.W., Hayhurst, J., Malangone, C., McMahon, A., Morales, J., Mountjoy, E., Sollis, E. et al. (2019) The NHGRI-EBI GWAS Catalog of published genome-wide association studies, targeted arrays and summary statistics 2019. Nucleic Acids Res, 47, D1005–D1012.

18. Magno, R. and Maia, A.T. (2020) gwasrapidd: an R package to query, download and wrangle GWAS catalog data. Bioinformatics, 36, 649–650.

19. Thul, P.J., Akesson, L., Wiking, M., Mahdessian, D., Geladaki, A., Ait Blal, H., Alm, T., Asplund, A., Bjork, L., Breckels, L.M. et al. (2017) A subcellular map of the human proteome. Science, 356.

20. Szklarczyk, D., Gable, A.L., Lyon, D., Junge, A., Wyder, S., Huerta-Cepas, J., Simonovic, M., Doncheva, N.T., Morris, J.H., Bork, P. et al. (2019) STRING v11: protein-protein association networks with increased coverage, supporting functional discovery in genome-wide experimental datasets. Nucleic Acids Res, 47, D607–D613.

21. Ruepp, A., Brauner, B., Dunger-Kaltenbach, I., Frishman, G., Montrone, C., Stransky, M., Waegele, B., Schmidt, T., Doudieu, O.N., Stumpflen, V. et al. (2008) CORUM: the comprehensive resource of mammalian protein complexes. Nucleic Acids Res, 36, D646–650.

22. Huttlin, E.L., Bruckner, R.J., Navarrete-Perea, J., Cannon, J.R., Baltier, K., Gebreab, F., Gygi, M.P., Thornock, A., Zarraga, G., Tam, S. et al. (2020) Dual Proteome-scale Networks Reveal Cell-specific Remodeling of the Human Interactome. bioRxiv, January 19, https://doi.org/10.1101/2020.1101.1119.905109.

23. Huttlin, E.L., Ting, L., Bruckner, R.J., Gebreab, F., Gygi, M.P., Szpyt, J., Tam, S., Zarraga, G., Colby, G., Baltier, K. et al. (2015) The BioPlex Network: A Systematic Exploration of the Human Interactome. Cell, 162, 425–440.

24. Eilbeck, K., Lewis, S.E., Mungall, C.J., Yandell, M., Stein, L., Durbin, R. and Ashburner, M. (2005) The Sequence Ontology: a tool for the unification of genome annotations. Genome Biol, 6, R44.

25. Hunt, S.E., McLaren, W., Gil, L., Thormann, A., Schuilenburg, H., Sheppard, D., Parton, A., Armean, I.M., Trevanion, S.J., Flicek, P. et al. (2018) Ensembl variation resources. Database (Oxford), 2018.

26. Cingolani, P., Platts, A., Wang le, L., Coon, M., Nguyen, T., Wang, L., Land, S.J., Lu, X. and Ruden, D.M. (2012) A program for annotating and predicting the effects of single nucleotide polymorphisms, SnpEff: SNPs in the genome of Drosophila melanogaster strain w1118; iso-2; iso-3. Fly (Austin), 6, 80–92.

27. Kustatscher, G., Grabowski, P., Schrader, T.A., Passmore, J.B., Schrader, M. and Rappsilber, J. (2019) Co-regulation map of the human proteome enables identification of protein functions. Nat Biotechnol, 37, 1361–1371.

28. R Core Team. (2017). R Foundation for Statistical Computing, Vienna, Austria.

29. Wickham, H., Averick, M., Bryan, J., Chang, W., McGowan, L., François, R., Grolemund, G., Hayes, A., Henry, L., Hester, J. et al. (2019) Welcome to the Tidyverse. Journal of Open Source Software, 4.

30. Gu, Z., Gu, L., Eils, R., Schlesner, M. and Brors, B. (2014) circlize Implements and enhances circular visualization in R. Bioinformatics, 30, 2811–2812.

31. Kang, H.W., Niepel, M.W., Han, S., Kawano, Y. and Cohen, D.E. (2012) Thioesterase superfamily member 2/acyl-CoA thioesterase 13 (Them2/Acot13) regulates hepatic lipid and glucose metabolism. FASEB J, 26, 2209–2221.

32. Gibbons, G.F. (2002) The role of cytochrome P450 in the regulation of cholesterol biosynthesis. Lipids, 37, 1163–1170.

33. Bartz, F., Kern, L., Erz, D., Zhu, M., Gilbert, D., Meinhof, T., Wirkner, U., Erfle, H., Muckenthaler, M., Pepperkok, R. et al. (2009) Identification of cholesterol-regulating genes by targeted RNAi screening. Cell Metab, 10, 63–75.

34. Sun, B.B., Maranville, J.C., Peters, J.E., Stacey, D., Staley, J.R., Blackshaw, J., Burgess, S., Jiang, T., Paige, E., Surendran, P. et al. (2018) Genomic atlas of the human plasma proteome. Nature, 558, 73–79.

35. Gold, B., Merriam, J.E., Zernant, J., Hancox, L.S., Taiber, A.J., Gehrs, K., Cramer, K., Neel, J., Bergeron, J., Barile, G.R. et al. (2006) Variation in factor B (BF) and complement component 2 (C2) genes is associated with age-related macular degeneration. Nat Genet, 38, 458–462.

36. Maller, J., George, S., Purcell, S., Fagerness, J., Altshuler, D., Daly, M.J. and Seddon, J.M. (2006) Common variation in three genes, including a noncoding variant in CFH, strongly influences risk of age-related macular degeneration. Nat Genet, 38, 1055–1059.

37. Catalan, V., Gomez-Ambrosi, J., Rodriguez, A., Ramirez, B., Rotellar, F., Valenti, V., Silva, C., Gil, M.J., Salvador, J. and Fruhbeck, G. (2012) Increased tenascin C and Toll-like receptor 4 levels in visceral adipose tissue as a link between inflammation and extracellular matrix remodeling in obesity. J Clin Endocrinol Metab, 97, E1880–1889.

38. Dickinson, M.E., Flenniken, A.M., Ji, X., Teboul, L., Wong, M.D., White, J.K., Meehan, T.F., Weninger, W.J., Westerberg, H., Adissu, H. et al. (2016) High-throughput discovery of novel developmental phenotypes. Nature, 537, 508–514.

39. Astle, W.J., Elding, H., Jiang, T., Allen, D., Ruklisa, D., Mann, A.L., Mead, D., Bouman, H., Riveros-Mckay, F., Kostadima, M.A. et al. (2016) The Allelic Landscape of Human Blood Cell Trait Variation and Links to Common Complex Disease. Cell, 167, 1415–1429 e1419.

40. McLachlan, S., Giambartolomei, C., White, J., Charoen, P., Wong, A., Finan, C., Engmann, J., Shah, T., Hersch, M., Podmore, C. et al. (2016) Replication and Characterization of Association between ABO SNPs and Red Blood Cell Traits by Meta-Analysis in Europeans. PLoS One, 11, e0156914.

41. Franchini, M., Capra, F., Targher, G., Montagnana, M. and Lippi, G. (2007) Relationship between ABO blood group and von Willebrand factor levels: from biology to clinical implications. Thromb J, 5, 14.

42. Yang, J., Li, W., Liu, S., Yuan, D., Guo, Y., Jia, C., Song, T. and Huang, C. (2016) Identification of novel serum peptide biomarkers for high-altitude adaptation: a comparative approach. Sci Rep, 6, 25489.

43. Emilsson, V., Ilkov, M., Lamb, J.R., Finkel, N., Gudmundsson, E.F., Pitts, R., Hoover, H., Gudmundsdottir, V., Horman, S.R., Aspelund, T. et al. (2018) Co-regulatory networks of human serum proteins link genetics to disease. Science, 361, 769–773.

44. Zhou, J., Wang, H., Lu, A., Hu, G., Luo, A., Ding, F., Zhang, J., Wang, X., Wu, M. and Liu, Z. (2002) A novel gene, NMES1, downregulated in human esophageal squamous cell carcinoma. Int J Cancer, 101, 311–316.

45. Court, M., Petre, G., Atifi, M.E. and Millet, A. (2017) Proteomic Signature Reveals Modulation of Human Macrophage Polarization and Functions Under Differing Environmental Oxygen Conditions. Mol Cell Proteomics, 16, 2153–2168.

46. Zimmer, A., Bouley, J., Le Mignon, M., Pliquet, E., Horiot, S., Turfkruyer, M., Baron-Bodo, V., Horak, F., Nony, E., Louise, A. et al. (2012) A regulatory dendritic cell signature correlates with the clinical efficacy of allergen-specific sublingual immunotherapy. J Allergy Clin Immunol, 129, 1020–1030.

47. Liu, G., Friggeri, A., Yang, Y., Park, Y.J., Tsuruta, Y. and Abraham, E. (2009) miR-147, a microRNA that is induced upon Toll-like receptor stimulation, regulates murine macrophage inflammatory responses. Proc Natl Acad Sci U S A, 106, 15819–15824.

48. Wang, L.J., Li, N.N., Xu, S.J., Zhang, F., Hao, M.H., Yang, X.J., Cai, X.H., Qiu, P.Y., Ji, H.L. and Xu, P. (2018) A new and important relationship between miRNA-147a and PDPK1 in radiotherapy. J Cell Biochem, 119, 3519–3527.

49. Preusse, M., Schughart, K. and Pessler, F. (2017) Host Genetic Background Strongly Affects Pulmonary microRNA Expression before and during Influenza A Virus Infection. Front Immunol, 8, 246.

